# Light-Activated Nucleic Acid Amplification Systems Using Photo-caged DNA Polymerase or Primers

**DOI:** 10.1101/2025.08.14.670248

**Authors:** Yang Wu, Meng Wu, Yukun Tian, Hao Chen, Xizhe Sun, Chukang Ma, Jiayao Liu, Fuyu Xia, Yu Liu, Xiaomeng Pei, Jing Wang, Guogang Zhao, Nan Wang, Fangfang Wang, Qing Liu, Fei Yu, Xiaofei Fan, Zehe Wang, Yue Chen, Jon West, Qi Cheng

## Abstract

Precise control over nucleic acid amplification is essential for achieving reliable, quantitative, high-throughput, and multiplexed molecular diagnostics, particularly in point-of-care and field settings. However, conventional isothermal amplification methods, especially rapid assays at ambient temperature such as recombinase-aided amplification (RAA), suffer from spontaneous initiation and lack of synchronization across parallel reactions. Here, we report light-activatable RAA systems employing two distinct photocaging strategies to achieve spatiotemporal control. In the first, a photocaged DNA polymerase was engineered via site-specific incorporation of p-azido-L-phenylalanine and conjugation with a 2-nitrobenzyl-modified ssDNA blocker through click chemistry. This approach offers high stability, making it advantageous for long-term storage and lyophilized formats. In the second, 5′-nitrobenzyl-modified primers were synthesized to block hybridization or extension until UV activation, offering operational simplicity and rapid integration into existing workflows. Both strategies independently suppressed background activity in the dark and restored full amplification efficiency upon 365 nm UV exposure. The systems exhibited high sensitivity, low background, and compatibility with portable diagnostics. A custom-built device integrating UV activation and real-time fluorescence detection enabled seamless, on-demand operation. This modular platform provides flexible light-gated amplification solutions, allowing the selection of polymerase- or primer-based caging according to storage stability or workflow requirements, thereby advancing precise and field-ready molecular diagnostics.

## 1 Introduction

Nucleic acid amplification technologies have revolutionized molecular diagnostics, enabling the rapid and sensitive detection of specific genetic material in fields ranging from clinical testing and food safety to environmental surveillance and synthetic biology^1,2^. While the polymerase chain reaction (PCR) remains the gold standard, its reliance on thermal cycling imposes constraints on instrumentation, energy requirements, and field deployment. As a result, isothermal amplification methods—such as loop-mediated isothermal amplification (LAMP), strand displacement amplification (SDA), and recombinase-aided amplification (RAA)—have attracted increasing attention for their ability to operate at constant temperatures using simpler equipment^1,3-5^.

Among these, recombinase-aided amplification (RAA) stands out due to its rapid reaction kinetics, low temperature requirement (typically 37–42°C), and compatibility with point-of-care testing (POCT)^6-8^. The RAA mechanism involves the coordinated action of a recombinase (UvsX), single-stranded DNA-binding protein (gp32), and strand-displacing DNA polymerase to facilitate primer invasion and target amplification^9-12^. Additionally, the inclusion of exonuclease III enables nicking of the DNA duplex, enhancing strand replacement and amplification efficiency^13,14^. With results achievable in under 30 minutes and detection limits in the low copy number range, RAA holds significant promise for decentralized molecular diagnostics^4,15-17^.

However, a critical limitation undermines the broader application and reliability of isothermal systems like RAA: lack of spatiotemporal control over reaction initiation^18,19^. In current formulations, the reaction begins spontaneously upon mixing components or upon addition of Mg^2^L ions, which complicates multiplexing, high-throughput workflows, and precise quantitative analyses^20^. Premature initiation during reaction setup or transport can lead to inconsistent results, false positives, and loss of synchronization among samples—problems that are exacerbated in resource-limited settings where environmental conditions are variable and manual handling is less controlled^21,22^.

To address this challenge, there is a growing interest in the development of externally controllable amplification systems, particularly those that can be activated by non-invasive and precise triggers^23-25^. Among various stimuli, light is particularly attractive due to its unique advantages: it offers fine spatial and temporal resolution, can be delivered remotely and rapidly, and does not require direct contact or chemical changes to the solution environment^26-28^. The use of photocaging strategies, in which biomolecular functions are reversibly inhibited by light-removable protecting groups, has opened new avenues in chemical biology, enabling optogenetic control of enzymes, transcription, and translation^29-31^. Extending this concept to nucleic acid amplification could yield a new generation of smart, programmable diagnostics.

Light-controlled strategies have emerged as a powerful means to achieve precise temporal and spatial regulation of nucleic acid amplification, addressing the challenges of background noise, premature initiation, and poor synchronization that often plague conventional amplification assays. In PCR and other isothermal methods such as LAMP, photoresponsive elements—most commonly nitrobenzyl derivatives—have been employed to cage key reaction components, including primers, nucleotides, and enzymes, thereby preventing unintended activity in the dark and enabling on-demand activation with ultraviolet light. These approaches offer advantages such as minimal chemical interference with reaction kinetics, reversible or irreversible control depending on the photocleavable group, and seamless integration with existing assay formats. Compared with systems that rely on thermal or chemical triggers, photoregulation allows non-invasive, rapid, and uniform activation across multiple parallel reactions. Given that recombinase-aided amplification (RAA) is highly sensitive yet prone to spontaneous initiation at room temperature, incorporating photocaging into the RAA workflow is both technically feasible and strategically advantageous. This combination leverages the intrinsic speed and simplicity of RAA with the precise controllability of photochemistry, enabling synchronized amplification, reduced false positives, and improved adaptability for point-of-care and field diagnostics.

In this study, we present the design and implementation of a light-activatable nucleic acid amplification system based on RAA, Pfu and Taq systems, using two photocaging strategies: one targeting the DNA polymerase, and another targeting the primers. For the enzyme-caging approach, we employed genetic code expansion and bioorthogonal chemistry to introduce a site-specific unnatural amino acid p-azido-L-phenylalanine (pAzF), into the DNA polymerase^32^. This azide handle was then conjugated to a single-stranded DNA (ssDNA) bearing a 2-nitrobenzyl photocleavable linker, effectively blocking the polymerase’s active site until exposed to UV light. Independently, we synthesized primers modified with a 5′-nitrobenzyl group, preventing hybridization or elongation until light-triggered cleavage occurred.

These photocaged components were incorporated into a fully reconstituted *in vitro* RAA system, which included the recombinase UvsX, loader UvsY, gp32, exonuclease III, and template DNA. Upon illumination at 365 nm, the system exhibited rapid and synchronized activation of amplification, while remaining completely inert under dark conditions. Both enzyme- and primer-level caging strategies were demonstrated to be effective, with excellent light-responsiveness, low background signal, and preservation of amplification fidelity and speed.

This photocontrol strategy represents a significant innovation in isothermal amplification technology. It offers several advantages: (1) precise temporal gating, ensuring consistent reaction start times; (2) elimination of spontaneous background amplification, improving assay specificity; (3) integration into portable, low-resource settings, where automation and hands-free operation are critical; and (4) potential for multiplexed or spatially patterned assays, enabled by the light of specific wavelengths.

Furthermore, we designed a customized device integrating ultraviolet light exposure and real-time fluorescence monitoring, enabling one-step light activation and detection. The platform is amenable to freeze-drying, making it suitable for field use and long-term storage.

In summary, this work introduces a modular, programmable, and field-ready nucleic acid detection system controlled by light. By combining the precision of photochemistry with the speed and simplicity of RAA, it lays the foundation for future development of intelligent biosensors, spatiotemporally regulated molecular tools, and next-generation diagnostics capable of operating in any environment—on demand, and with confidence.

## 2 Results

### 2.1 RAA Construction and Optimization

To establish a reliable platform for isothermal nucleic acid amplification, a complete recombinase-aided amplification (RAA) system was constructed using recombinantly expressed and purified protein components. The essential proteins—recombinase UvsX, recombinase loader UvsY, single-stranded DNA-binding protein gp32, DNA polymerase (Sau), and exonuclease III (Exo)—were individually cloned and expressed in *E*.*coli*. SDS-PAGE analysis confirmed the successful expression of all target proteins, with distinct bands corresponding to their expected molecular weights (Figure S1). Following affinity purification via Ni-NTA chromatography, the proteins exhibited high purity and were suitable for downstream reconstitution of the RAA system (Figure S2).

Agarose gel electrophoresis was employed initially to assess the functionality of the *in vitro* assembled RAA system. In the presence of all five core protein components, robust bands were observed within 30 minutes at 37□°C, indicating that the system was catalytically active and properly reconstituted (Figure 1A). A fluorescence-based amplification assay targeting a specific DNA sequence was also employed (Figure 1B)

**Figure 1.**
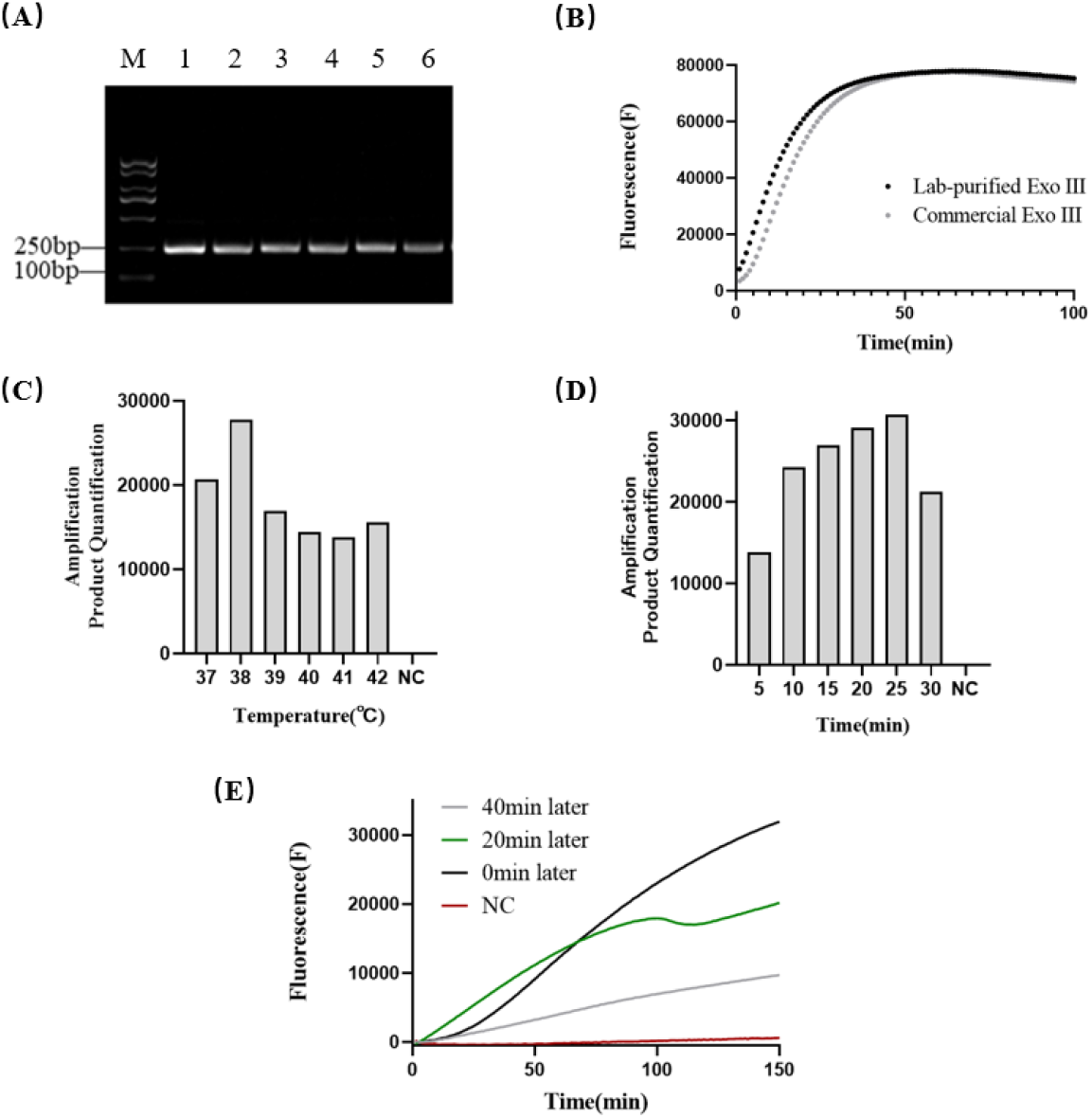
RAA construction and optimization. **(A)** Functional reconstitution of the RAA system using laboratory-prepared components. Lane 1: commercial RAA kit as a positive control; Lanes 2–5: individual replacement of DNA polymerase, recombinase (UvsX), co-recombinase (UvsY), or single-stranded DNA-binding protein (gp32) with in-house purified enzymes; Lane 6: complete replacement of all components. Amplification of the 177 bp target fragment was successfully observed in all lanes, confirming the activity and compatibility of the reconstituted system. **(B)** Comparison of amplification performance using commercial versus lab-purified exonuclease III (Exo III) in a fluorescence-based RAA assay. Both variants produced comparable signal intensity, verifying the functional equivalence of the in-house enzyme. **(C)** Optimization of reaction temperature. Amplification efficiency at various temperatures (e.g., 33– 42□°C) was assessed by gel electrophoresis followed by grayscale quantification. The optimal performance was observed at 38□°C. **(D)** Optimization of reaction time. Grayscale analysis of agarose gel electrophoresis results at different time points (e.g., 5–30 min) revealed that 25 minutes provided the highest amplification efficiency and was selected as the optimal reaction duration. **(E)** Evaluation of synchronization in conventional RAA reactions. Sequential preparation of multiple RAA reaction tubes under room temperature conditions led to variable amplification kinetics, suggesting spontaneous, asynchronous initiation among replicates.

Subsequently, key reaction parameters were optimized to enhance performance. The amplification efficiency was evaluated across a temperature gradient (30°C to 45°C). The optimal reaction temperature was identified as 38°C, consistent with previously reported RAA systems (Figure 1C). Reaction kinetics were further examined over time to determine the shortest effective amplification window. Fluorescence monitoring revealed that product accumulation plateaued within 25 minutes under optimal conditions, supporting a practical diagnostic timeframe (Figure 1D).

Despite the system’s high efficiency under optimized conditions, a critical limitation was observed during delayed reaction setup. When the reaction mixture was incubated at room temperature before addition of magnesium, significant variability in amplification kinetics was noted across parallel samples. Some replicates initiated amplification prematurely, while others showed delayed or inconsistent signal onset (Figure 1E). This stochastic behavior indicates that the current RAA formulation lacks strict temporal control and is prone to spontaneous reaction initiation during long preparation. This drawback highlights the urgent need for an externally controllable activation mechanism to synchronize reaction initiation—an issue addressed by the light-responsive strategy developed in the subsequent sections.

### 2.2 Construction and Characterization of Photocaged DNA Polymerase

To achieve light-dependent control of DNA polymerase activity, a chemically modified enzyme was engineered via site-specific incorporation of an unnatural amino acid. The target enzyme, derived from Staphylococcus aureus (Sau) DNA polymerase, was genetically modified to include an amber (TAG) codon. This codon allowed incorporation of the photo-reactive amino acid p-azido-L-phenylalanine (pAzF) via an orthogonal tRNA/aminoacyl-tRNA synthetase pair in a recoded *E. coli* C321.ΔA.exp strain. After induction with IPTG in the presence of pAzF, the modified enzyme was expressed and purified using nickel affinity chromatography.

SDS-PAGE analysis (Figure 2A) confirmed successful overexpression of the full-length unnatural DNA polymerase, showing expected molecular weight by mass spectrum (Figure 2B), and no significant protein degradation was observed. Compared to the wild-type enzyme, the pAzF-incorporated polymerase retained high expression yield and solubility, supporting its suitability for subsequent chemical modification and functional testing.

**Figure 2.**
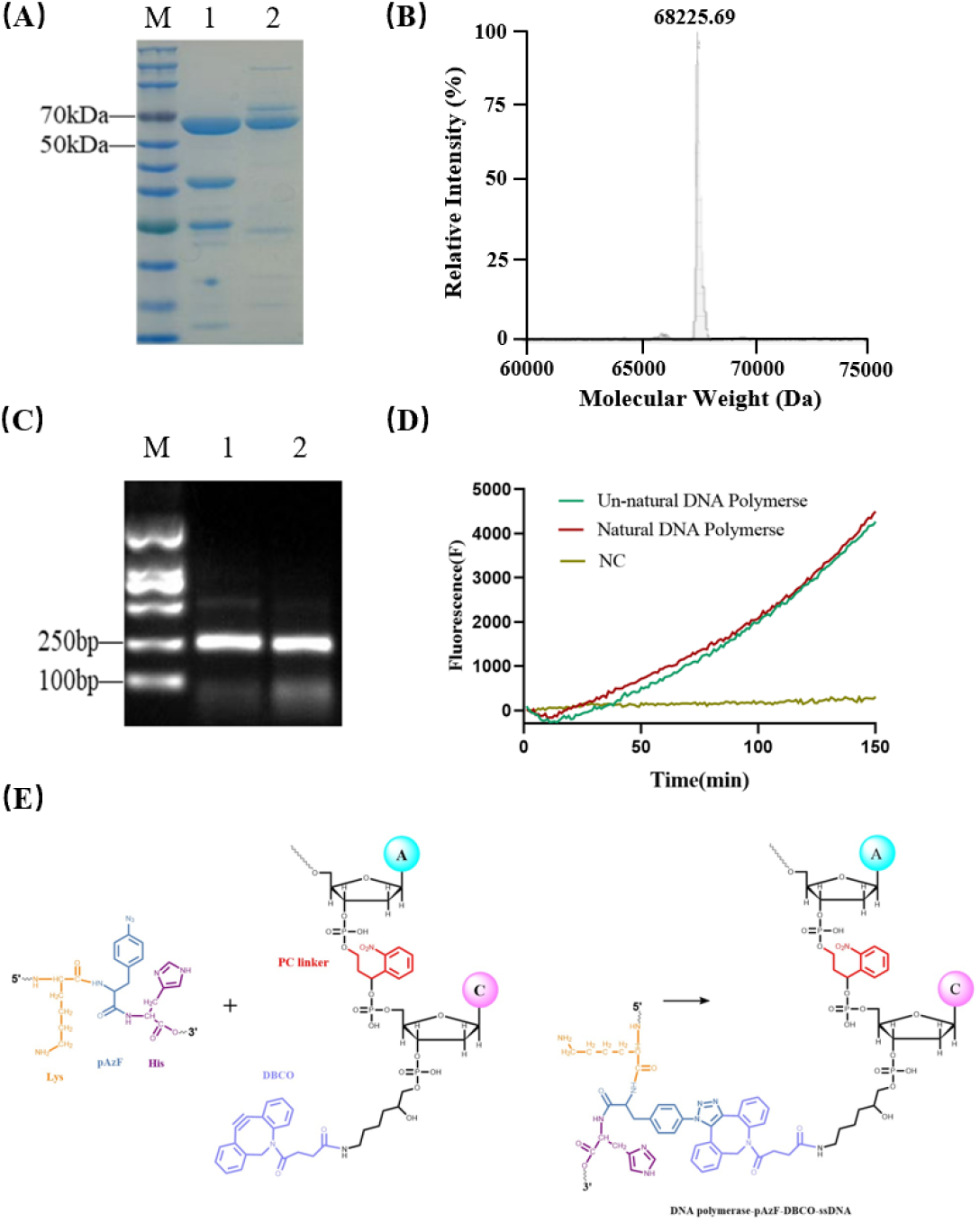
Construction and Characterization of Photocaged DNA Polymerase. **(A)** SDS-PAGE analysis comparing wild-type and unnatural DNA polymerase. Lane 1: wild-type polymerase; Lane 2: polymerase containing the site-specific incorporation of the unnatural amino acid p-azido-L-phenylalanine (pAzF). Both proteins show a clear band at approximately 68.4 kDa, indicating successful expression and similar purity. **(B)** Molecular weight determination of the unnatural DNA polymerase using liquid chromatography– mass spectrometry (LC-MS). The mass spectrum reveals a single major peak corresponding to 68,225.69 Da, consistent with the expected mass of the full-length protein containing pAzF. **(C)** Gel-based activity assay of the unnatural DNA polymerase in the RAA system. Replacement of the wild-type polymerase with the unnatural variant (lane 2) produced comparable amplification of the target band to that observed with the native enzyme (lane 1), demonstrating retained catalytic activity. **(D)** Fluorescence-based RAA assay confirming the enzymatic functionality of the unnatural DNA polymerase. Fluorescence amplification kinetics using the unnatural enzyme mirrored those of the wild-type, further validating its compatibility with RAA. **(E)** Schematic representation of the strain-promoted azide–alkyne cycloaddition (SPAAC) reaction between the azide-functionalized unnatural DNA polymerase and a dibenzocyclooctyne (DBCO)-modified, nitrobenzyl photocleavable single-stranded oligonucleotide, forming the photocaged polymerase via click chemistry.

To evaluate whether the unnatural amino acid substitution altered enzymatic activity, the polymerase was subjected to in vitro RAA assays using both agarose gel electrophoresis and real-time fluorescence quantification. As shown by the agarose gel results (Figure 2C), the unnatural DNA polymerase retained DNA synthesis activity similar to the wild-type enzyme in the absence of any chemical modification, indicating that the pAzF incorporation did not intrinsically disrupt polymerase function. Complementary fluorescence-based kinetic analysis further confirmed the ability of the enzyme to support efficient amplification of target nucleic acids, with fluorescence intensities and reaction rates comparable to the native enzyme (Figure 2D).

To achieve light-responsiveness, the azide group on the incorporated pAzF residue was conjugated with a DBCO-modified single-stranded DNA (ssDNA) containing a 2-nitrobenzyl photocleavable moiety (Figure 2E and Figure S3). This bio-orthogonal “click” reaction (strain-promoted azide-alkyne cycloaddition, SPAAC) covalently tethered the ssDNA to the enzyme, spatially blocking its catalytic site and thereby inhibiting its polymerase activity in the absence of light. The efficiency of this conjugation was confirmed by mobility shift assays and the absence of amplification signal in dark conditions.

This chemically caged polymerase served as a central component of the light-activatable nucleic acid amplification platform. Upon exposure to 365 nm UV light, the photocleavable linker was rapidly cleaved, releasing the blocking ssDNA fragment and restoring enzymatic activity. The successful design and functional verification of the photocaged polymerase established a robust molecular switch for temporal control of DNA amplification reactions, forming the foundation for a light-regulated isothermal system described in the following sections.

### 2.3 Light-Activated RAA System

Following the successful construction of a photocaged DNA polymerase, the next objective was to establish a functional light-activated recombinase-aided amplification (RAA) system that could be precisely triggered by external light input. A critical parameter influencing the success of polymerase caging was the molar ratio of the azide-containing enzyme to the DBCO-modified ssDNA blocker. The ssDNA was designed to sterically hinder the polymerase active site upon conjugation via strain-promoted azide-alkyne cycloaddition (SPAAC). To determine the optimal stoichiometry, a series of conjugation reactions were performed with varying ssDNA-to-polymerase ratios, and the resulting conjugates were tested for background amplification in the absence of light.

Quantitative analysis demonstrated that polymerase activity decreased progressively with increasing ssDNA concentration, plateauing at a molar ratio of approximately 2.5:1 (Figure 3A). Beyond this point, no significant improvement in inhibition was observed, indicating saturation of the available reactive azide sites. This ratio was chosen for subsequent experiments to ensure maximum suppression of activity without excess unbound ssDNA, which could otherwise interfere with the recombinase or template interactions. This step was crucial for ensuring the enzyme remained fully inactive under dark conditions until deliberately uncaged.

**Figure 3.**
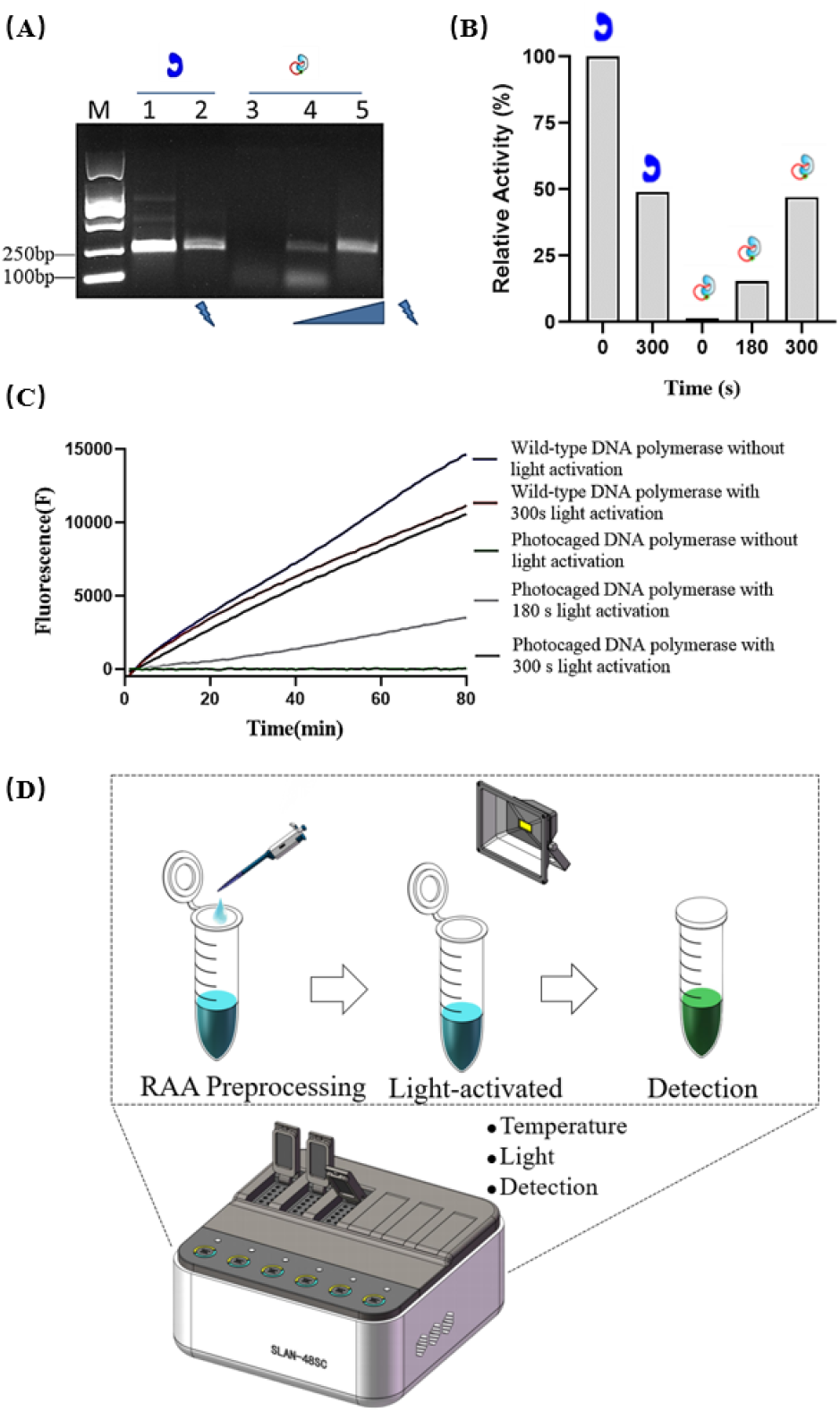
Light-Activated RAA System. **(A)** Agarose gel electrophoresis analysis of DNA amplification using photo-caged DNA polymerase under different light activation conditions. Lane 1: wild-type DNA polymerase without UV activation (positive control); Lane 2: wild-type polymerase after 300 s UV activation; Lane 3: photo-caged DNA polymerase without UV exposure, showing no amplification; Lane 4: photo-caged polymerase after 180 s UV activation, showing weak amplification bands; Lane 5: photo-caged polymerase after 300 s UV activation, showing clear and strong amplification, indicating successful light-induced enzyme activation. **(B)** Grayscale quantification of gel bands shown in panel A. Band intensities were measured and plotted as bar graphs corresponding to each lane, confirming that polymerase activity and amplification efficiency increase proportionally with UV exposure time. **(C)** Fluorescence-based analysis of the light-activated RAA system. The same sample setup as in panel A was tested in real-time fluorescence assays. Only UV-activated reactions with photo-caged polymerase produced significant fluorescence signals, demonstrating successful amplification and system responsiveness to light. **(D)** Schematic diagram of an integrated light-activated amplification device. The platform combines 365 nm UV illumination with fluorescence detection and temperature control in a compact design, enabling synchronized activation and real-time monitoring of nucleic acid amplification for field-deployable diagnostics.

After determining the optimal conjugation conditions, a complete light-controlled RAA system was assembled by integrating the photocaged polymerase with recombinase proteins (UvsX and UvsY), gp32, exonuclease III, primers, template, and necessary ions. The mixture was incubated under light-protected conditions to prevent premature activation. Upon exposure to 365 nm ultraviolet light for 180 seconds at 20 watts, the photocleavable linker in the ssDNA was cleaved, releasing the steric blockade and restoring the catalytic activity of the DNA polymerase.

The effectiveness of light-triggered amplification was validated using multiple analytical methods. Agarose gel electrophoresis clearly demonstrated that DNA amplification only occurred in samples exposed to UV light. As shown in Figure 3A, irradiated samples produced strong, specific DNA bands, whereas samples kept in the dark showed no visible amplification product, confirming the strict light dependency of the system. Grayscale analysis of the gel bands (Figure 3B) further quantified the amplification efficiency, with signal intensities correlating directly with light exposure and confirming that the system remained effectively inert until triggered.

To further quantify the amplification process, real-time fluorescence monitoring was employed. Amplification kinetics revealed a sharp increase in fluorescence intensity immediately following UV illumination, as shown in Figure 3C. In contrast, fluorescence signals in the dark control group remained at baseline throughout the reaction course. These results confirmed that the photocaged RAA system enables both suppression of unwanted background activity and rapid activation upon light exposure, resolving a major limitation of traditional isothermal amplification methods.

To facilitate practical application and improve usability, an integrated reaction platform was developed to combine both light activation and fluorescence-based detection in a compact format. The device comprises a programmable 365 nm LED illumination unit, a temperature-controlled reaction chamber, and an integrated optical sensor for real-time fluorescence readout. The schematic in Figure 3D illustrates the system’s ability to perform both initiation and monitoring steps seamlessly, thereby enabling synchronized activation across multiple reaction chambers. This design supports potential applications in point-of-care diagnostics and high-throughput screening, offering a controllable and automatable nucleic acid detection workflow.

In summary, these results confirm the successful construction of a light-responsive RAA platform with tight spatiotemporal control. The system eliminates spontaneous background amplification, supports precise initiation timing, and is compatible with field-deployable detection formats, laying the groundwork for further development of programmable nucleic acid amplification tools.

### 2.4 Light-Activated Nucleic Acid Amplification Systems by Photocaged Primers

To complement the photocaged DNA polymerase strategy and broaden the versatility of light-regulated nucleic acid amplification, we next focused on the development of photocaged primers capable of independently controlling reaction initiation. This primer-caging approach introduces a photoremovable group at a strategic position on the oligonucleotide, blocking primer-template hybridization or extension until cleaved by ultraviolet light. Such a method offers operational simplicity and modular design, and is compatible with multiple polymerase systems, including RAA, Taq, and Pfu.

A series of primers were chemically modified with a 2-nitrobenzyl photocleavable group, a well-characterized moiety that is stable under ambient conditions but rapidly cleaved upon irradiation at 365 nm. As summarized in Table 1, both forward and reverse primers targeting a specific amplicon sequence were synthesized in caged and uncaged formats. The chemical modification was introduced during solid-phase oligonucleotide synthesis, and the final products were purified by high-performance liquid chromatography (HPLC). Mass spectrometry analysis confirmed the successful incorporation of the protecting groups, with precise molecular weights matching theoretical values for each primer.

**Table 1.**
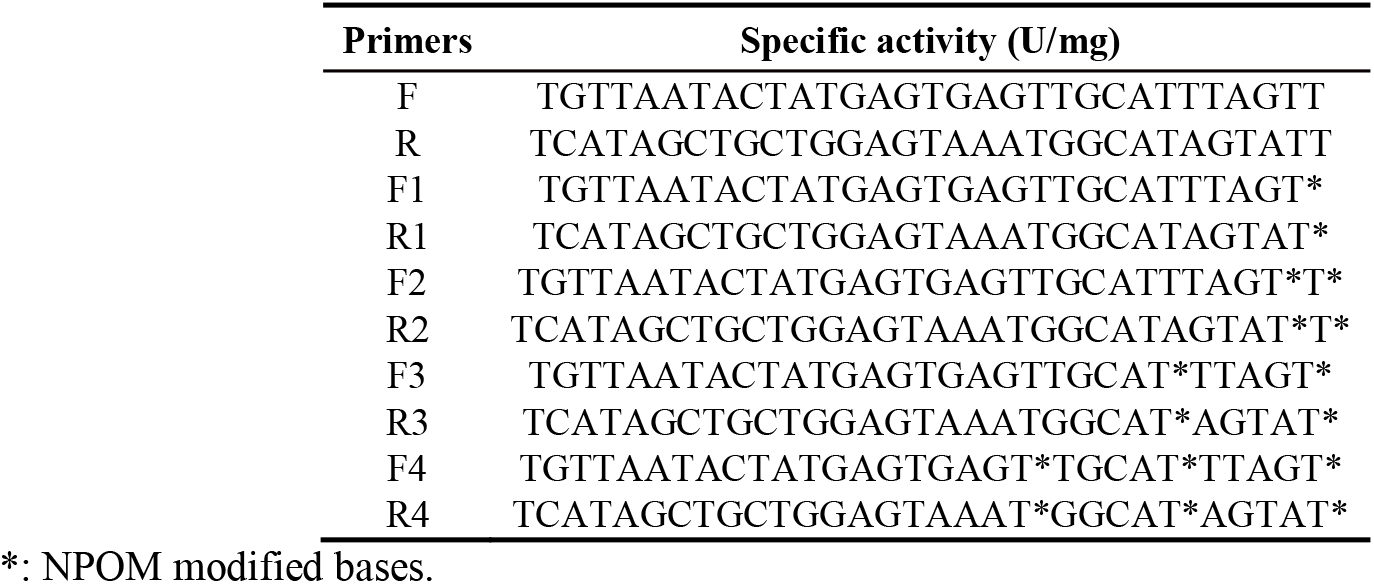
The number and position of NPOM Caged-dT modifications in primers.

To assess the functional integrity of the caged primers, activity assays were conducted using three different amplification systems: the isothermal RAA platform, and two representative thermocycling enzymes—Taq polymerase and Pfu polymerase. In each case, amplification reactions were prepared with the same core components, substituting only the forward or reverse primer with its caged variant. Reactions were carried out under standard conditions and divided into two groups: those exposed to 365 nm UV light for 180 seconds and those maintained in the dark.

For the RAA system (Figure 4C), results showed that caged primers completely suppressed nucleic acid amplification in the absence of UV light, as confirmed by both agarose gel electrophoresis and fluorescence signal tracking. Upon light exposure, strong amplification signals were restored, closely matching those observed with non-caged primers. These findings confirmed that the photocaging group effectively blocked recombinase-mediated strand invasion and polymerase extension, while remaining fully reversible upon illumination.

**Figure 4.**
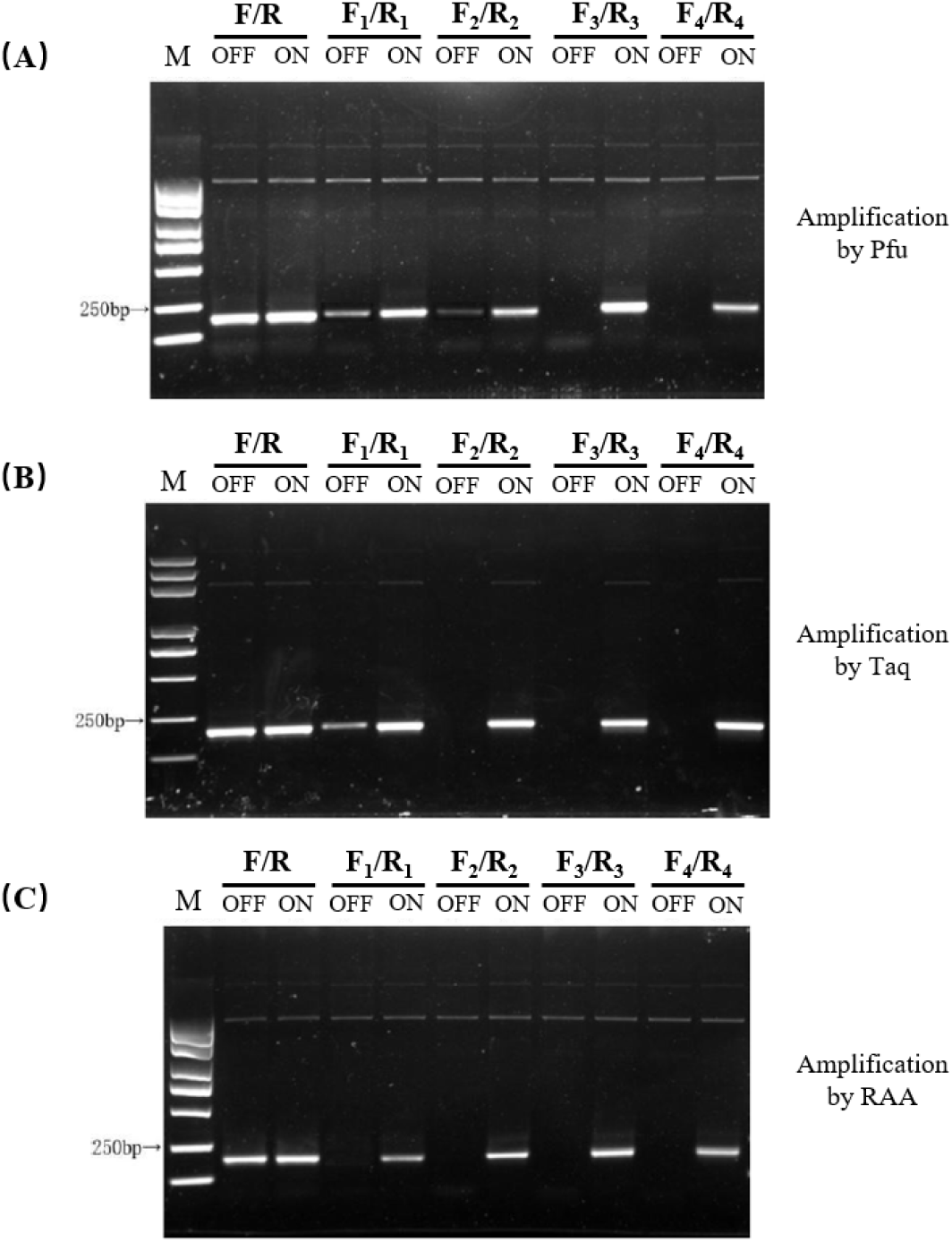
Light-Activated Nucleic Acid Amplification Systems by Photocaged Primers. Photocaged primers designed as listed in Table 1 were evaluated in three different nucleic acid amplification systems: Pfu polymerase **(A)**, Taq polymerase **(B)**, and RAA **(C)**. For each system, reactions were performed under two conditions: without UV light activation (**OFF** lanes) and with 365 nm UV light activation (**ON** lanes).

A similar trend was observed in Taq polymerase-mediated PCR (Figure 4B). Amplification reactions containing caged primers failed to generate any visible DNA bands in the dark, indicating successful inhibition of primer function. Post-irradiation reactions, however, yielded clear and specific amplification products, confirming the complete restoration of primer activity. These results suggest that the bulky nitrobenzyl group likely obstructs primer annealing or polymerase binding at the primer-template junction, and that light-induced cleavage of the group is sufficient to recover full enzymatic function.

In the case of Pfu polymerase (Figure 4A), which possesses 3′ to 5′ exonuclease proofreading activity, the photocaged primers similarly exhibited a complete suppression of amplification under dark conditions, without any detectable degradation or partial extension. Light activation restored activity with comparable efficiency to standard primers, highlighting the robustness of the caging strategy across different enzyme classes.

Together, these experiments demonstrate the universality and functional reliability of photocaged primers across both isothermal and thermocycling amplification platforms. Their ability to provide strict light-dependent control over primer function—regardless of polymerase type—establishes them as a powerful tool for programmable molecular diagnostics. This modular photocaging approach not only supports temporal regulation but also opens possibilities for multiplexed, spatially-resolved amplification schemes and integration into field-deployable diagnostic platforms.

## 3 Discussion

In this study, we developed and systematically validated a novel light-activatable nucleic acid amplification system based on photocaged DNA polymerase and photocaged primers into a recombinase-aided amplification (RAA) platform. This optical control architecture enables precise spatiotemporal regulation over the initiation of isothermal DNA amplification reactions, addressing one of the critical limitations of conventional isothermal methods: spontaneous, asynchronous background activity during sample preparation.

The implementation of photocaged DNA polymerase was achieved through the site-specific incorporation of an unnatural amino acid, p-azido-L-phenylalanine (pAzF), into the polymerase backbone, followed by covalent conjugation with a 2-nitrobenzyl-modified single-stranded DNA blocker via bioorthogonal click chemistry. This strategy effectively rendered the enzyme catalytically inert under ambient conditions and reactivated it upon UV exposure. Parallel to this, photocaged primers were synthesized with a 5′-nitrobenzyl protecting group that prevented hybridization or elongation until removed by light. Both caging mechanisms independently exhibited complete suppression of amplification under dark conditions and rapid, full restoration of activity after brief UV light activation. These properties were validated across multiple assay platforms—including agarose gel electrophoresis, fluorescence quantification, and grayscale imaging—and tested with different enzymes (RAA polymerase, Taq, and Pfu), establishing the broad applicability and orthogonality of the approach.

Compared to traditional magnesium-dependent initiation methods used in commercial RAA kits, this light-activatable system provides a significant advancement by offering synchronized activation across multiple reactions, even in long-term or high-throughput workflows. The capacity to precisely control reaction onset using light avoids the stochastic behavior caused by temperature fluctuations or inconsistent magnesium chelation kinetics. This level of control is especially advantageous in point-of-care testing (POCT) settings, where sample processing and reaction setup may occur under variable environmental conditions.

Moreover, the modularity and tunability of both caging strategies are noteworthy. The location, chemistry, and number of photocaging groups can be modified to fine-tune the level of inhibition and activation kinetics. This flexibility allows the design of multi-stage, orthogonally addressable amplification systems activated by different wavelengths of light or triggered in a programmable sequence. Such innovations could open the door to complex nucleic acid computing systems or spatially resolved biosensing platforms.

From a practical perspective, the integration of this dual light-activatable system into a miniaturized, field-compatible reaction device further enhances its translational value. The custom-built platform combines 365 nm UV irradiation with real-time fluorescence detection in a compact, automation-friendly format, making it suitable for field diagnostics, high-throughput screening, or synthetic biology applications. Importantly, the freeze-drying compatibility of the system also supports ambient storage and transportation without compromising stability or responsiveness.

Despite these promising results, several limitations remain. First, the use of UV light for activation, while effective, may pose concerns regarding potential DNA damage and safety in live-cell or clinical environments. Future studies may explore red-shifted photocaging groups that respond to visible or near-infrared light, thereby mitigating phototoxicity and improving tissue penetration for potential in vivo applications. Second, although the current system achieves efficient on-demand activation, it primarily operates as an endpoint trigger. The capability for dynamic modulation of amplification speed, output levels, or feedback regulation remains unexplored. Incorporating optical logic gates, photoswitchable molecular beacons, or light-responsive inhibitors could enhance the system’s programmability and functional diversity. Third, from a translational perspective, the synthesis of photocaged oligonucleotides and the preparation of photocaged enzymes currently require specialized reagents and multi-step protocols, which may increase production costs relative to conventional amplification reagents. Reducing synthetic complexity or developing scalable, low-cost photocaging chemistries will be critical for broader adoption, particularly in resource-limited diagnostic settings. Addressing these challenges will be essential to fully realize the potential of light-gated RAA systems in both laboratory research and practical point-of-care applications.

In conclusion, this work presents a versatile and robust light-regulated nucleic acid amplification platform that incorporates chemically engineered photocaged DNA polymerases and primers. It provides precise temporal control, low background amplification, and compatibility with both isothermal and PCR workflows. This dual-caging architecture lays a strong foundation for future development of intelligent, programmable nucleic acid detection systems and has broad implications in diagnostics, synthetic biology, environmental monitoring, and biosensing. Future directions will focus on expanding light responsiveness to longer wavelengths, achieving multiplexed control, and integrating the system into microfluidic and wearable diagnostic devices for next-generation molecular diagnostics.

## 4 Materials and methods

### 4.1 Construction and Purification of RAA Core Components

The core protein components required for the RAA system were recombinantly expressed in *E. coli* and purified using standard affinity chromatography. The target genes for DNA polymerase (e.g., Sau or Pfu), recombinase (UvsX), auxiliary protein (UvsY), single-stranded DNA binding protein (gp32), and exonuclease III (Exo) were cloned into expression vectors such as pET28a or pTrc99A. Protein expression was induced with 1 mM IPTG at 16–20°C, and cell pellets were lysed by sonication. The resulting supernatants were subjected to Ni-NTA affinity purification, followed by dialysis into reaction-compatible buffers. Protein concentrations were determined via BCA assay, and purity was confirmed by SDS-PAGE.

### 4.2 Engineering of Photocaged DNA Polymerase

To develop a light-activatable DNA polymerase, site-specific incorporation of the unnatural amino acid p-azido-L-phenylalanine (pAzF) into the polymerase sequence was performed. A TAG codon was introduced to the polymerase using overlap extension PCR. Expression was carried out in *E. coli* C321.ΔA.exp strains containing an orthogonal tRNA/tRNA synthetase pair specific for pAzF. Induction conditions and pAzF concentration were optimized for maximal yield and incorporation efficiency.

Following protein purification, the azide group of the incorporated pAzF was reacted with a DBCO-modified DNA oligonucleotide bearing a 2-nitrobenzyl photocleavable group. This bioorthogonal click reaction (SPAAC) resulted in a sterically hindered polymerase that could not initiate strand elongation until exposed to 365 nm light. The success of conjugation and restoration of activity after illumination were verified by gel electrophoresis and enzymatic assays.

### 4.3 Synthesis and Characterization of Photocaged Primers

Forward and reverse primers were chemically modified with a photocleavable 2-nitrobenzyl group, which prevents hybridization or elongation. These photocaged primers were synthesized commercially, purified by HPLC. Primer stability and uncaging efficiency under 365 nm light were evaluated *in vitro*.

### 4.4 Photocontrol and Functional Validation

Enzyme and primer photoactivation were characterized independently. Polymerase activity was measured by adding a fluorescently labeled dsDNA template and quantifying extension using real-time fluorescence. Primer functionality was assessed by their ability to initiate amplification with an uncaged polymerase. Time-course UV illumination experiments (30 to 300 seconds) were conducted to determine optimal activation windows. Control groups included no-light, no-primer, and uncaged conditions to assess background signal.

### 4.5 Light-Activated RAA Amplification System

The complete RAA system was reconstituted using purified UvsX, UvsY, gp32, Exo, and either the photocaged or native polymerase, along with the light-activatable primers. Reactions were set up at 37°C and shielded from ambient light. UV activation was achieved using a 365 nm LED panel (manufactured by Foshan Lighting Factory) positioned 25 cm from the samples, operated at 20 watts for 180–300 s, followed by real-time fluorescence monitoring (excitation 495 nm, emission 520 nm) for nucleic acid amplification detection. The fluorescence threshold time (Tt) and reaction kinetics were analyzed using a qPCR instrument or portable POCT device.

### 4.6 Lyophilization and Field-Compatible Format

To enhance portability and storage, the complete system was lyophilized with trehalose and PEG as stabilizers. The lyophilized reagents were vacuum-sealed under dry conditions and stored at room temperature. Prior to use, reagents were rehydrated with Mg^2^□ solution and irradiated with UV light to initiate the reaction.

## 5 CRediT authorship contribution statement

QC conceived the presented idea and supervised this work. MW, YKT and HC conducted the major experiments. YW, XZS, CKM, YL, XMP, NW, JW, FFW, QL, FY guided the experiments and drafted the original manuscript. All authors have read and approved the final version of the manuscript.

## 6 Acknowledgments

This work was supported by the National Science Foundation of China (Grant No. 32471477 to QC), Hebei Natural Science Foundation (Grant No. C2024204170 to YW) and the Talents Introduction Plan of Hebei Agricultural University (Grant No. YJ2023025 to YW).

---

**Figure S1.**
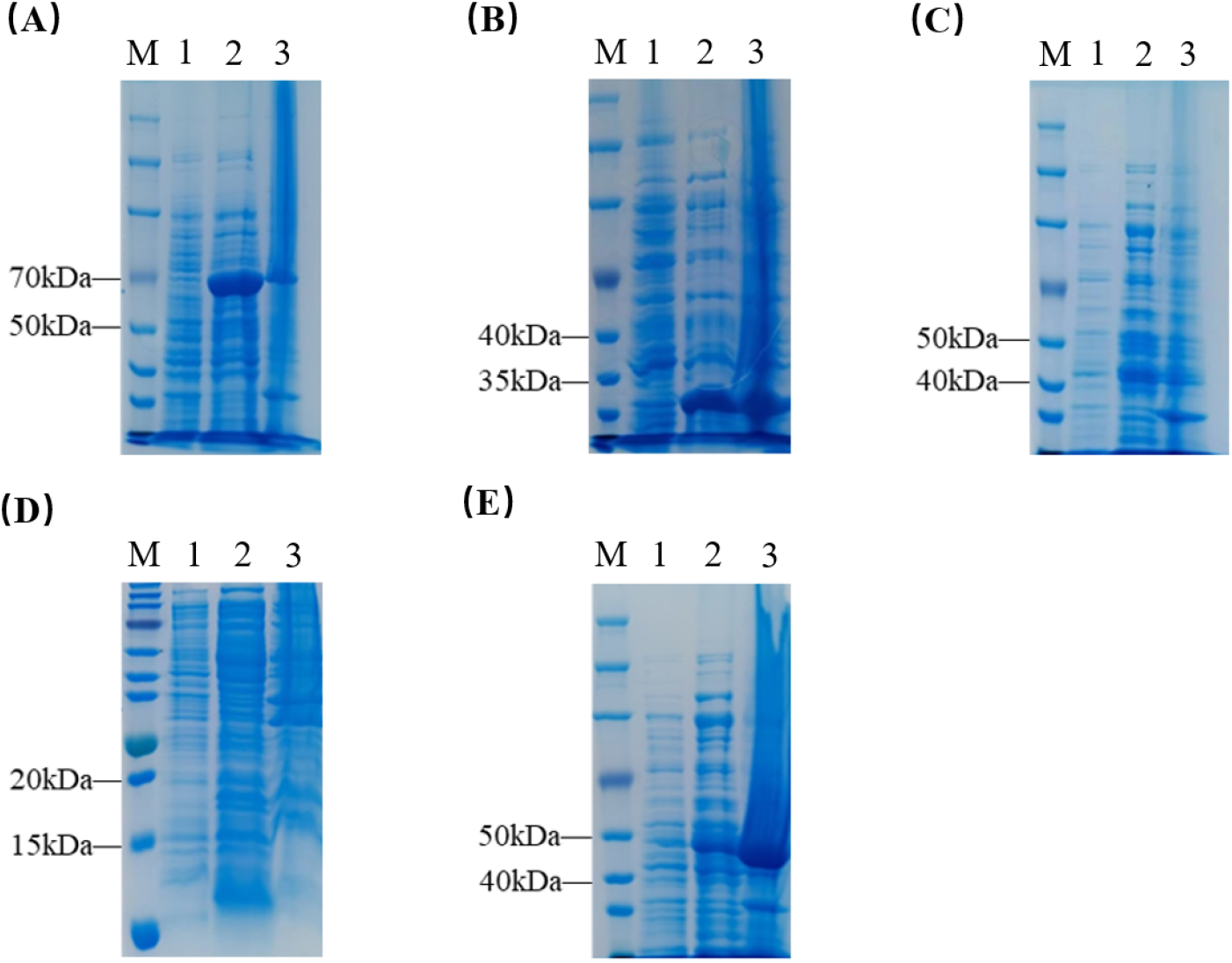
SDS-PAGE analysis of protein expression for five key RAA components in *E. coli*. (A) DNA polymerase; (B) single-stranded DNA-binding protein (gp32); (C) recombinase (UvsX); (D) co-recombinase (UvsY); (E) exonuclease III (Exo III). For each protein, Lane 1: total lysate before IPTG induction; Lane 2: soluble fraction after IPTG induction; Lane 3: insoluble fraction (inclusion bodies) after IPTG induction.

**Figure S2.**
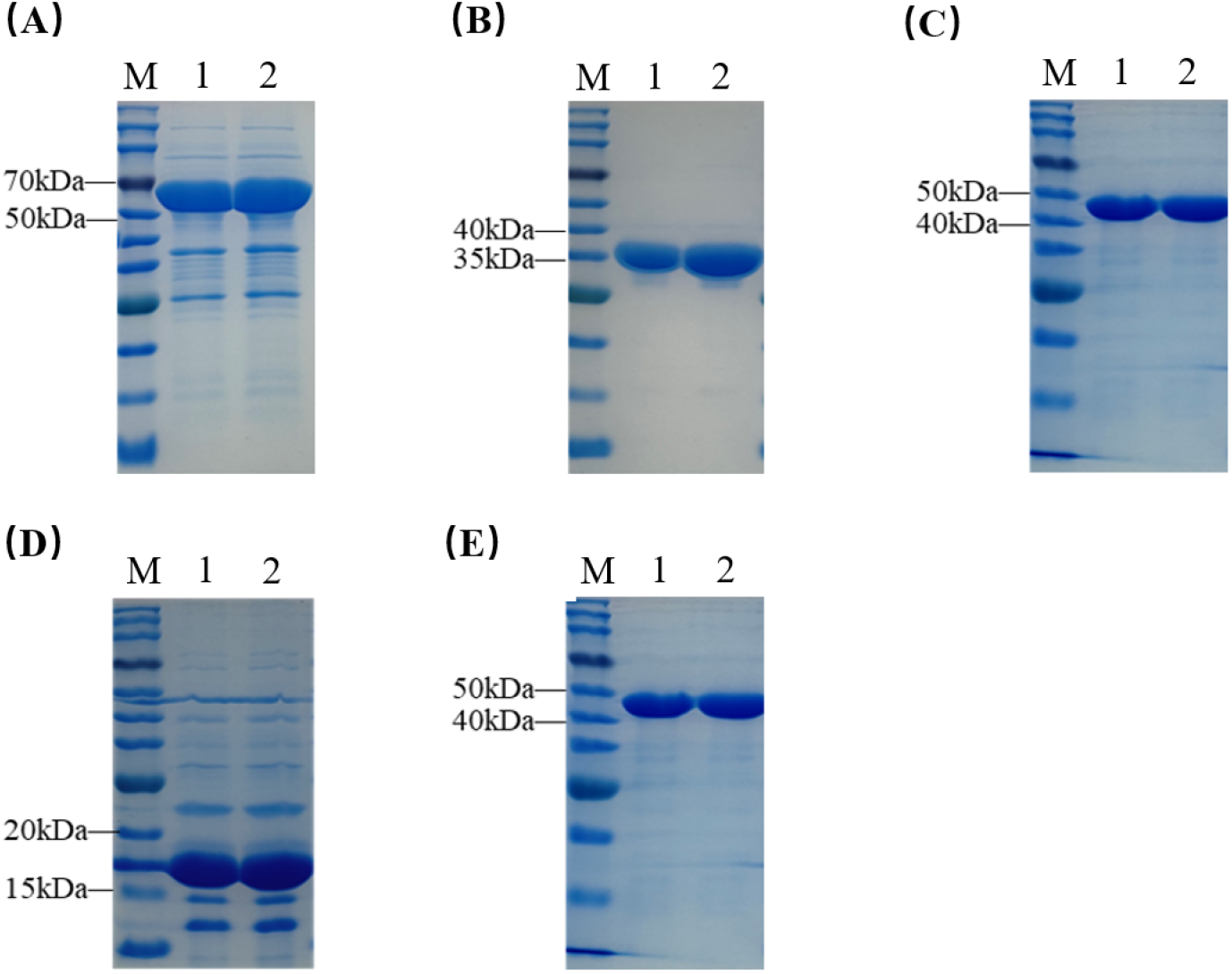
SDS-PAGE analysis of purified recombinant proteins used in RAA system. (A) DNA polymerase; (B) single-stranded DNA-binding protein (gp32); (C) recombinase (UvsX); (D) co-recombinase (UvsY); (E) exonuclease III (Exo III). For each protein, Lane 1: purified protein sample before dialysis; Lane 2: protein sample after dialysis.

**Figure S3.**
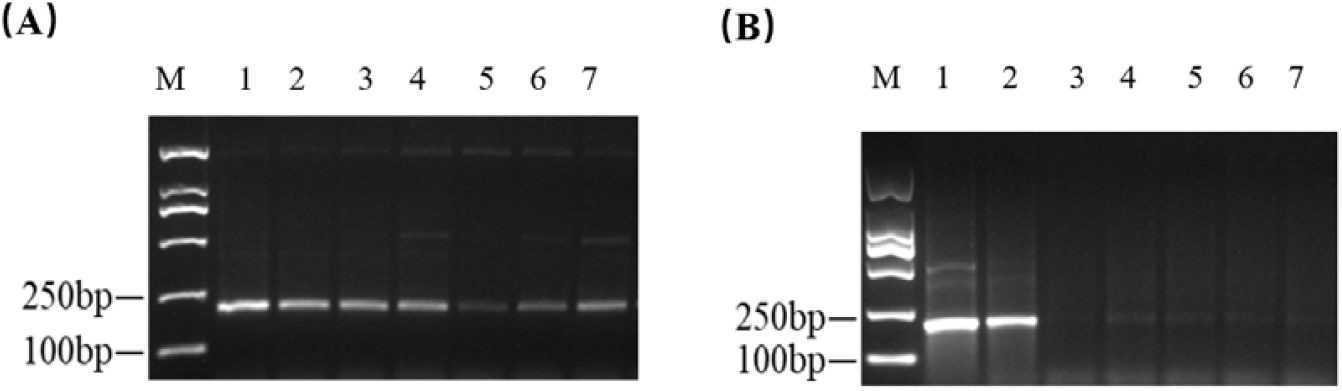
Optimization of the Conjugation Ratio Between Azide-Containing DNA Polymerase and DBCO-Modified Single-Stranded DNA (ssDNA) Blocker. Assessment of the optimal molar ratio for the strain-promoted azide–alkyne cycloaddition (SPAAC) reaction between azide-functionalized DNA polymerase and DBCO-modified ssDNA blocker used to generate photocaged polymerase. **(A)** Conjugation performed at a 1:2 molar ratio (enzyme:ssDNA); **(B)** Conjugation performed at a 1:3 molar ratio. After conjugation, the activity of the resulting photocaged polymerase was evaluated in the RAA system with and without UV light activation. In both panels, **Lane 1–2**: amplification products using wild-type (uncaged) DNA polymerase as positive controls; **Lane 3**: photocaged polymerase without UV irradiation, showing suppressed amplification; **Lanes 4–5**: photocaged polymerase after 180 seconds of UV activation, showing partial restoration of amplification; **Lanes 6–7**: photocaged polymerase after 300 seconds of UV activation, demonstrating enhanced recovery of enzymatic activity.

